# Importance of the Landlocked Pond Smelt *Hypomesus nipponensis* as a food resource of the Little Tern *Sternula albifrons* on inland Andong Lake of Korea: A video image analysis

**DOI:** 10.1101/660142

**Authors:** Dong-Man Shin, Jeong-Ho Han

## Abstract

We carried out the diet study of the little tern on the sandy islet in inland Andong Lake, Korea, during the beeding season (April to July 2018). To identify its diet and examine the importance of the main prey species as a food resource, we set two remote-control video cameras with 4K-resolution on the islet. One thousand two hundred seventy-five still images that the tern had prey in its bill were identified at the species level and measured on a monitor. Then, they were classified to five length-categories and compared among months and breeding stages. Freshwater fishes dominated the diet (100%; eleven species overall), where the landlocked pond smelt *Hypomesus nipponensis* (80.8%) and largemouth bass *Micropterus salmoides* (13.7%) were the primary and next essential prey species, respectively. The average prey item length was 51.04 ± 20.89 mm and significantly differed among months and breeding stages (*P* < 0.001, respectively). 50–75 mm prey length category was the most frequent in the diet (42.2%). In April and May, larger fish >50 mm constituted the greater part of their diet (93.1%, 66.3%, respectively), whereas the diet in June and July consisted of smaller fishes <50 mm (56.2%, 68.8% respectively). The occurrence frequency of prey length categories also varied significantly among the breeding stages (*P* < 0.001): 1–25 mm and 50–75 mm were overrepresented and underrepresented, respectively, at the chicks in the nest stage. On the other hand, 50–75 mm was preferred for the pre-laying and incubation stages. In terms of the survival condition of pond smelts, the before- and after water surface temperatures of the day when terns flew away showed a significant difference (*P* = 0.004), where a threshold looks like between 29.11°C and 30.04°C. These results support the prey abundance hypothesis that, when cold-water pond smelts might wholly swim down into the deeper lake in the hot summer, the terns might also leave their colony for another foraging place with higher prey availability.

## Introduction

The little tern is a seabird predator that breeds on spits of sand, shingle, and shell fragments on seashores or in estuaries as well as rivers, lakes, and reservoirs [1, 2]. It preys mostly on small fishes and invertebrates, mainly insects and crustaceans [3]. To date, however, many studies have focused on its breeding, diet, foraging, predation, and habitat degradation on offshores or in estuaries [4–8], which might result from scarce information in inland lakes [9].

Over the past 30 years, little terns had formed their breeding colonies in Nakdong Estuary (4,759 individuals) [10], Ganwol Lake (516 nests) [11], and Sihwa Reclaimed Area (300 pairs) [12], which had been their main habitats in Korea. However, due to the destruction and disturbance of natural habitats, predation, and vegetation, they have no longer visited these breeding sites before 4 or 5 years [13, 14]. In the meanwhile, it turned out that a part of little terns bred on an islet in inland Andong Lake (for detail, see Study area), about 170 km away northern from Nakdong Estuary, since 2005 [15, 16]. Nevertheless, a more in-depth study on their breeding ecology and diet has been scarcely carried out in this habitat. Based on discarded preys on the ground, it was simply reported that they foraged on pond smelts (*Hypomesus nipponensis*) [15].

There are two types of pond smelts in Korea: a resident one which grows and spawns in the lake, and an anadromous migration one which migrates downstream to the sea and returns to the lake or inflowing river for spawning [17–19]. They commonly prefer to swim and prey on zooplankton near the water surface below 20°C, whereras they swim down in deeper water when the water temperature increases in hot summer [20, 21]. For Andong Lake, they are resident fishes, transplanted in 1981 for enlarging a catch of fish [20–22], as was in the Korean other lakes since 1925 [23, 20]. Bird’s diet in any particular location is usually dominated by one or two species, typically those that are locally abundant and profitable [24, 25]. The little tern, an opportunistic forager [26], depends on the abundance of the main prey, and thus its diet reflects prey availability in the habitat. If the seabird would prey on the landlocked pond smelt abundant in a freshwater lake, it means that the cold-water fish [20] could inhabit nearer water surface during tern’s breeding season from April to July, particularly in hot summer. It could probably play a leading prey role such as herring, sandeel [27], sand-smelt (*Atherina* spp.), goby (*Pomatoschistus* spp.) [26], dusky trident goby (*Tridentiger obscurus*) [5], pilchard (*Sardinops neopilchardus*), southern anchovy (*Engraulis australis*), and sprat (*Spratelloides robustus*) [28], which are its main food resources in beach or estuary habitats.

There have ever been numerous methods to investigate the diet of piscivorous birds [29–31]: (1) to collect otoliths and scales around nest [32], (2) collect discarded preys on the ground [5, 33], (3) induce regurgitation [34], (4) photograph birds carrying preys [35, 36] and so on. Despite of many merits and effectiveness, these diet studies have each limitation that it may be possible not to reveal the entire diet of the bird or to be invasive. In this regards, the remote-control video camera (hereafter RCVC) deserves much consideration. The latest ones can record images at the quality of ultra high definition (4K), making bird researchers gain diverse information on diets since they offer enlarged but vivid images profitable for prey identification. Video shooting far away from the breeding colony can also prevent field researchers from interfering in bird’s incubating or feeding activities and give them more information in real-time or on SD card if needed.

The objectives of this study were (1) to identify the little tern diet and determine its main prey species (probably pond smelt) on the images taken by 4K RCVC, (2) examine its survival condition, and (3) discuss the importance as a food resource. We expect that it could contribute to the protection and conservation of the unique seabird habitat located on an inland lake.

## Methods

### Study area

The diet study was carried out on a sandy islet of Andong Lake (36° 34′ 35.06″ N, 128° 49′ 53.98″ E; Fig 1), Andong City, located on inland Korea, during the breeding season from April to July in 2018. This Lake, the second largest one in Korea (after Soyang Lake), was made by Andong Dam which was completed in 1976 to supply agricultural and industrial water and electric power, with controlling flood [16]. It has a water storage area of 51.5 Km^2^ and high water level of 160 EL m. Its last 10-year average water level and precipitation are 144.86 EL m and 1080.56 mm, respectively [37], and the average temperature is 18.83°C during the breeding season and particularly 24.3°C (max: 29.0°C) in July [38].

**Fig 1.**
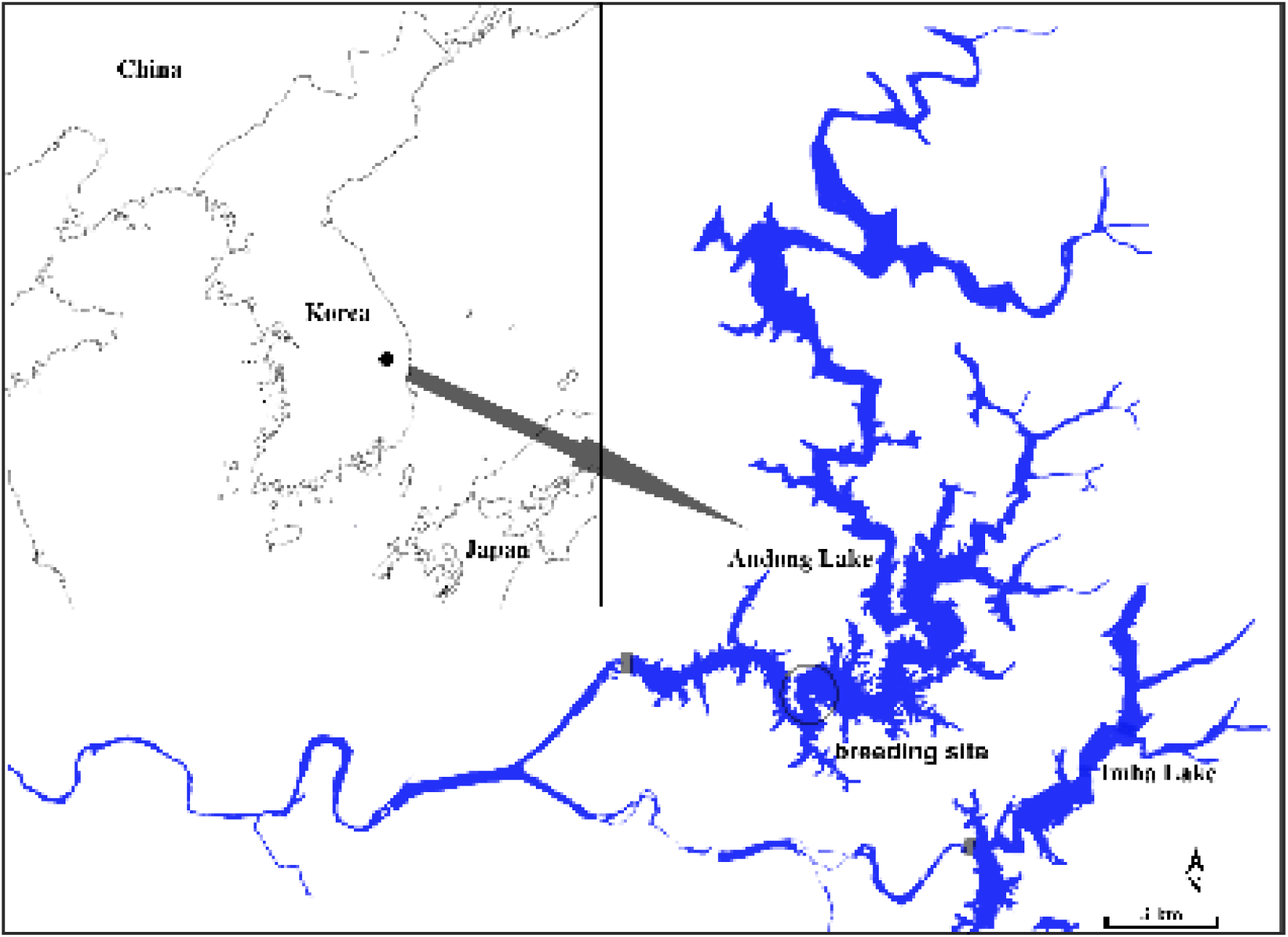
Andong Lake located in inland Korea, where the diet of little terns was studied, April to July 2018.

The islet on the lake where little terns bred consisted of two parts, sandy (146.66 EL m) and rocky (146.93 EL m), where the one occupied the majority. Its area was approximately 100 × 20 × 4 m in length, width, and height during the breeding season, which could irregularly change according to water level [Fig 2]. However, there was found a certain periodicity every year [37, Fig 3]: It submerged entirely under the water after it rained heavily in summer, with appearing again in spring. In general, the depth of water around the islet was about 20–30 m. Ichthyofauna in the lake consisted mainly of Cyprinidae, Centrarchidae, and Osmeridae [16].

**Fig 2.**
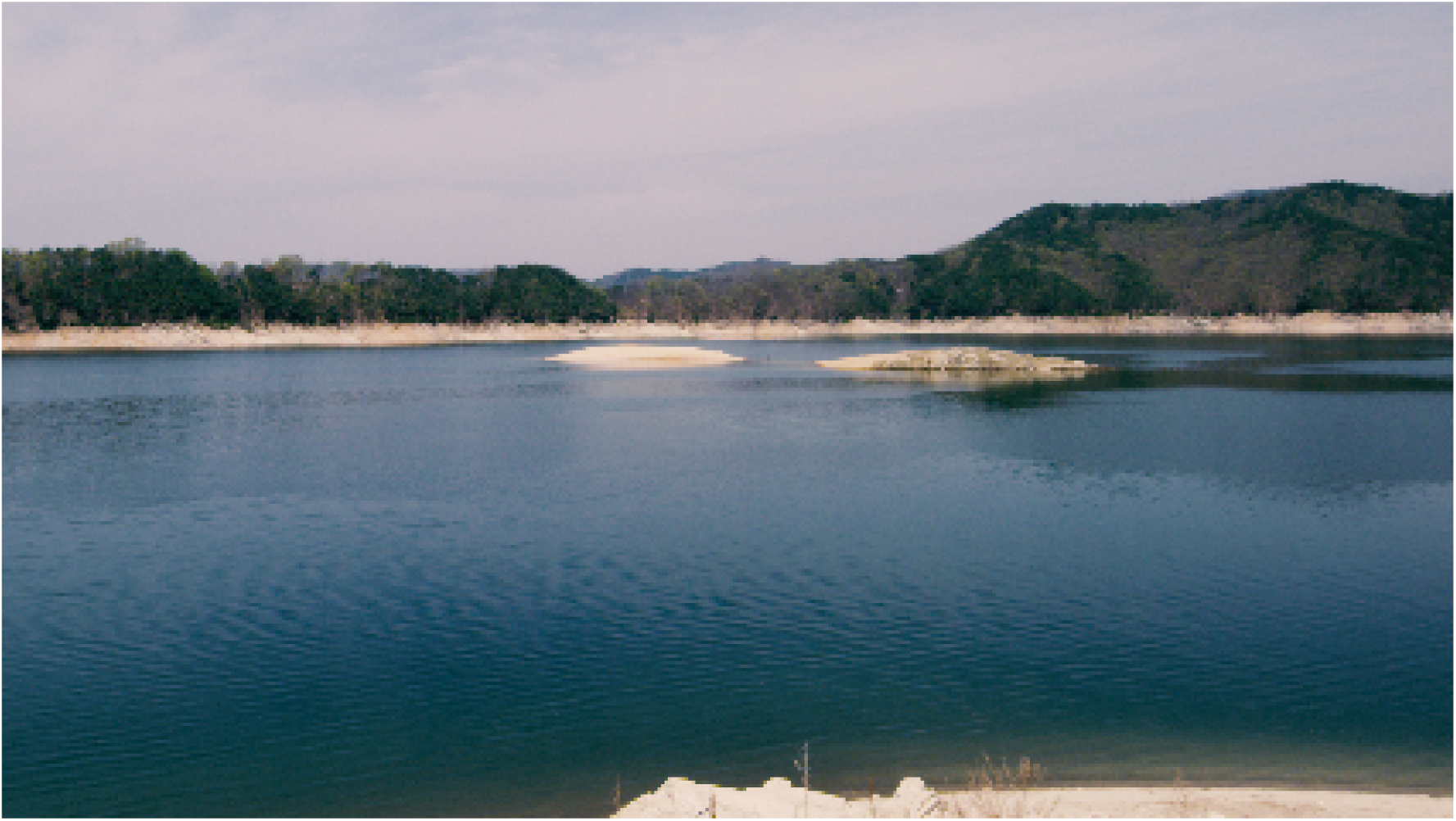
Islet that little terns bred, located on inland Andong Lake, Korea.

**Fig 3.**
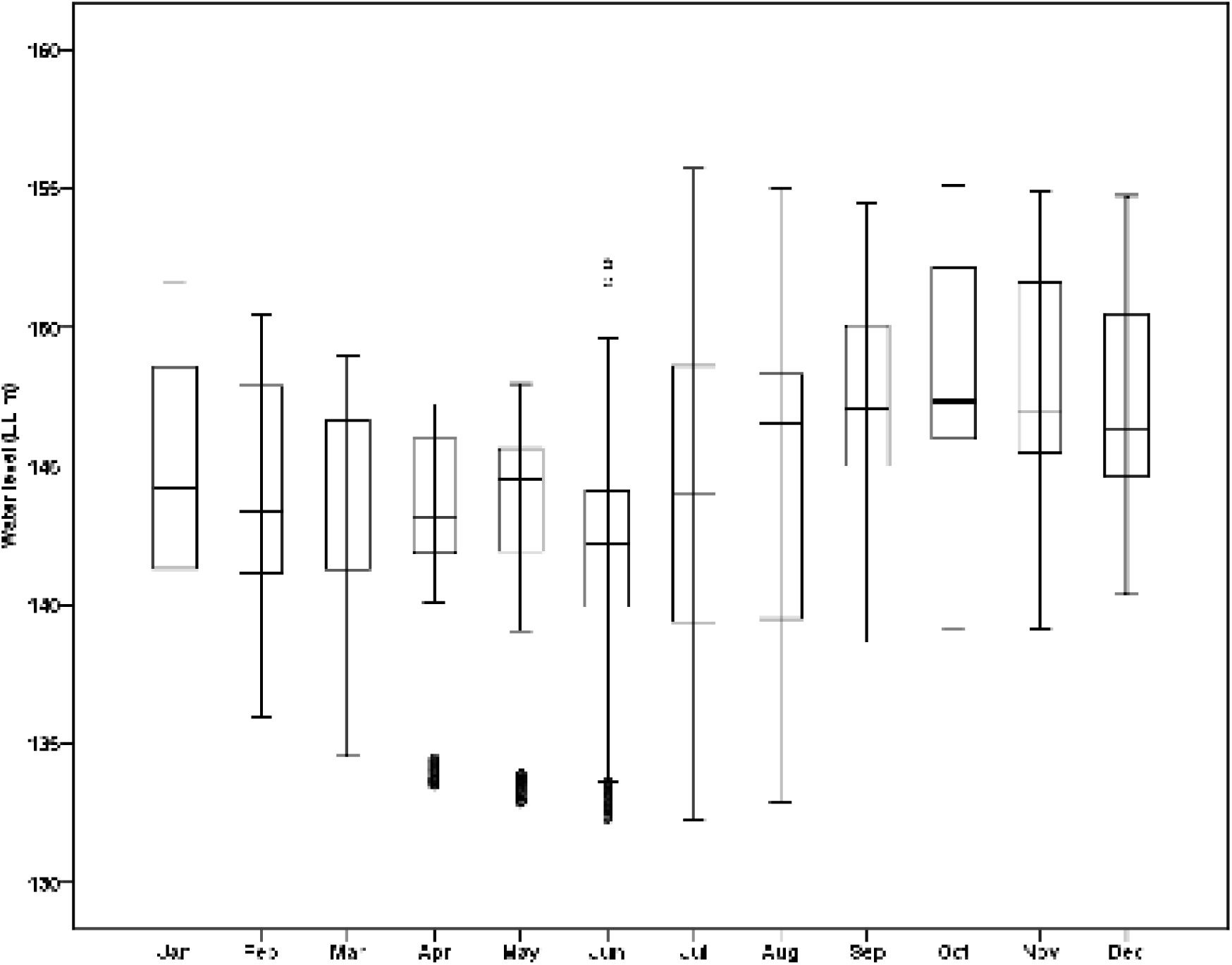
Water level (EL m) per month between 2008 and 2017 on inland Andong Lake, Korea.

### Image-sampling

To take pictures of prey items that little terns carried in their bills, we set 2 remote-control video cameras [DS-2DF883615X-AEL(W), Hikvision; hereafter RCVC] with an one m-high pole at both high point and corner of the islet. Camera installation was conducted in late March before the terns arrived, not to disturb their breeding activity. We built a temporary barrack on the land 350 m away from the islet under permission of the public authority and recorded video files in there by network video recorder (DS-7716NI-I4, Hikvision) that could receive video signals using WiFi signals. Since the video camera had a function of IR mode at night, it was possible to take around-the-clock shooting. Besides, we could focus, zoom (36×), and pan (360°) camera remotely. Main image specification was as follows: resolution (3840×2160), frame rate per second (30), and image compression (H.265). Except for special days, i.e., checking a bad WiFi signal and temporarily withdrawing camera due to an increase in water level, we usually recorded tern’s prey delivery events 17 h a day from 05:00 to 22:00. After chicks left the islet, we shot parent birds’ feeding behaviors by a portable 4K-video camera (Raven, RED).

### Identification of prey species

To identify prey items, we captured their still images using a professional video-editing package (Premiere 2018, Apple). To avoid duplicate and multiple calculations, we selected one image per prey delivery event. Based on local fish literature [39–41], we identified prey at the level of species and determined its numerical abundance [42]. Then, we classified undistinguishable long shots or blurred pictures into “unidentified”. The frequency of prey occurrence was compared among months (April–July) and breeding stages: (1) the pre-laying (Male brings prey to female for mating), (2) the incubation (Male delivers prey to incubating female), (3) the chicks in the nest (Parents deliver prey to chicks in the nest), and (4) the mobile chicks on the ground period (Parents deliver prey to frisking chicks).

### Calculation of prey length

To obtain prey length data, we also measured little tern’s culmen length (LCL) and prey’s total length (PTL) on a monitor (resolution: 1920 × 1080, length: 15.6 inches) by a vernier caliper (nearest 0.01 mm). Then, we selected only side shots or frontal shots with a vertical bill because those with a diagonal bill to the camera had become logarithmically shorter in LCL than their actual values [43]. In addition, we excluded those in which even an end of bill or prey was covered with other subjects in measurement for minimizing bias. The little tern carries fish crosswise in its bill [44], being hung vertically, where there may be less distorted in length. Since a head of the fish, however, tends to tilt to some extent, we measured the PTL in consideration of the unfolded length of the bent fish after a preliminary measurement. Although the male tern is greater in body size than the female, it is not easy to visually distinguish between the two sexes [45]. Therefore, we determined the actual PTL by comparing the measured LCL and PTL and average LCL (30.56 mm) measured in Korea [11].

With a division of 25 mm, we complementarily classified PTLs to 5 length categories: (1) 1–25 mm, (2) 25–50 mm, (3) 50–75 mm, (4) 75–100 mm, (5) ≥100 mm. Based on this classification information, we determined the preferred prey size of little tern adults and chicks as follows, respectively: (1) very small, (2) small, (3) medium, (4) large, and (5) very large. To verify temporal variation in prey size, we also analyzed the frequency of these categories in 4 months and breeding stages, respectively.

### Aquatic factors

The investigation of the water surface temperature and dissolved oxygen (hereafter WST and DO, respectively) was conducted to assess their effects on tern’s primary prey. Since the tern opts to prey mainly on fish nearer the surface or in water less than 15 cm deep [46, 28, 47], we surveyed the water factors between the surface and 1 m deep during the breeding season. We obtained these data from automatic measurement system (Troll 9500, Insitu) set by dam management authority, which automatically measured them four times a day (06:00, 12:00, 18:00, and 23:00). We used the average value of both depth categories and 4-hour categories as a day value. The WST and DO of the day at which all little terns migrated into different foraging place in the late breeding season were defined as potential survival thresholds, under which the main prey fish could no longer survive and thus moved into deeper water. By comparing both “before-” and “after-” moving values (1st to 15th and 16th to 31th, 15th and 16th of July, respectively) the correlations between aquatic factors, little terns, and pond smelts were examined.

### Data analysis and Statistics

Chi-squared test and Kruskal-Willis test were used to assess differences in diet composition and occurrence frequency of prey length categories and ones in average PTL between months and breeding stages, respectively. However, a comparison of two length variables or “before-” and “after-” moving” values was conducted by Mann-Whitney *U*-test or Student *t*-test. We expressed data as mean ± SD and set the significance level at *P* < 0.05 for all statistical tests. All computations were carried out in PASW Statistics 18.

## Results

The date when little terns arrived for the first time at the sand islet on Andong Lake was the 9th of April, and the one when they left for the last time was the 16th of July. In total, 31 breeding nests were made there, and 78 eggs were laid and incubated. We gained video clips of ca. 2,000 h 89 days long and succeeded to capture 1,275 best still images that terns had preys in their bills.

Most of the images (97.2%) were identified at the species level. Freshwater fishes dominated the diet, accounting for 100% of prey by number and a total of 11 species (Table 1). On the contrary, insects and crustaceans did not occur in the current study. For identified prey items, the pond smelt (80.8%) was the primary prey species, and the largemouth bass (*Micropterus salmoides*; 13.7%) was the next essencial one (Table 1). The latter appeared suddenly on the prey identification list in late May, which consisted mostly of very small and small fries (for bass 47.4%, 33.7%, respectively; Table 1 and Fig 4). The other nine fish species constituted a minor part of the diet (2.8%; Table 1).

**Table 1.** Diet composition of little terns breeding on a sandy islet in Andong Lake of inland Korea.

| Identified prey items | n | F (%) |
| --- | --- | --- |
| **Osmeridae** | **1,030** | **80.8** |
| *Hypomesus nipponensis* | 1,030 | 80.8 |
| **Centrarchidae** | **175** | **13.7** |
| *Micropterus salmoides* | 175 | 13.7 |
| **Cyprinidae** | **33** | **2.8** |
| *Erythroculter erythropterus* | 20 | 1.6 |
| *Hemiculter eigenmanni* | 3 | 0.2 |
| *Opsarichthys u. amurensis* | 2 | 0.2 |
| *Squalidus chankaensis tsuchigae* | 2 | 0.2 |
| *Hemibarbus labeo* | 2 | 0.2 |
| *Pseudogobio esocinus* | 1 | 0.1 |
| *Pungtungia herzi* | 1 | 0.1 |
| *Zacco koreanus* | 1 | 0.1 |
| *Zacco platypus* | 1 | 0.1 |
| Unidentified | 37 | 2.9 |
| Total | 1,230 | 100.0 |

Prey species was analyzed on the images taken by 4K remote-controlled video cameras.
Number of each species (n) and frequency of each species (F) in the diet are shown.

**Fig 4.**
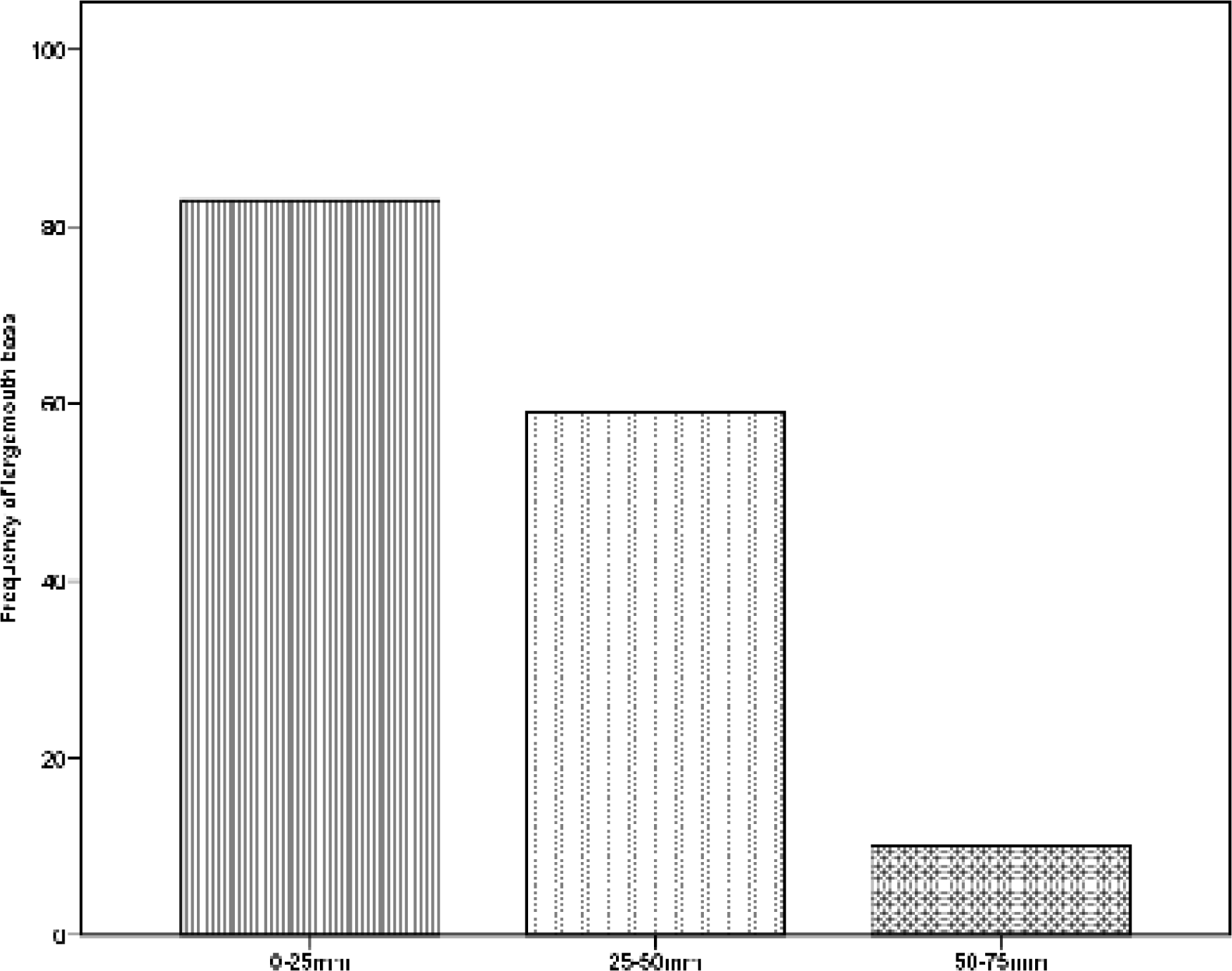
Occurrence frequency of the largemouth bass on the little tern prey images, compared between prey length categories.

The average prey item length was 51.04 ± 20.89 mm (n = 981, range = 11.23–112.93 mm) and significantly differed among both months and breeding stages (Kruskal-Willis Test: n = 981, χ^2^ = 104.624, *P* < 0.001; n = 979, χ^2^ = 213.243, *P* < 0.001, respectively; Fig 5A and 5B). In particular, 50–75 mm prey length category was the most frequent in the diet (42.2%), which means that here terns foraged mainly on medium-size fishes, and 25–50 mm one was the next common (31.4%). The average prey length tended to decrease according to months. In April and May, early breeding season, larger fish >50 mm constituted the greater part of the little tern diet (93.1%, n = 87, mean = 68.18 ± 13.20 mm, range = 36.66–105.81 mm; 66.3%, n = 279, mean = 54.59 ± 19.12 mm, range = 11.23–106.08, respectively; Fig 6A). On the other hand, the diet in June and July, chicks-rearing period, consisted of smaller fishes <50 mm (56.2%, n = 555, mean = 47.25 ± 21.36 mm, range = 11.97–112.93 mm; 68.8%, n = 60, mean = 44.66 ± 17.94 mm, range = 20.12–105.49 mm, respectively; Fig 6A and 6B).

**Fig 5A, 5B.**
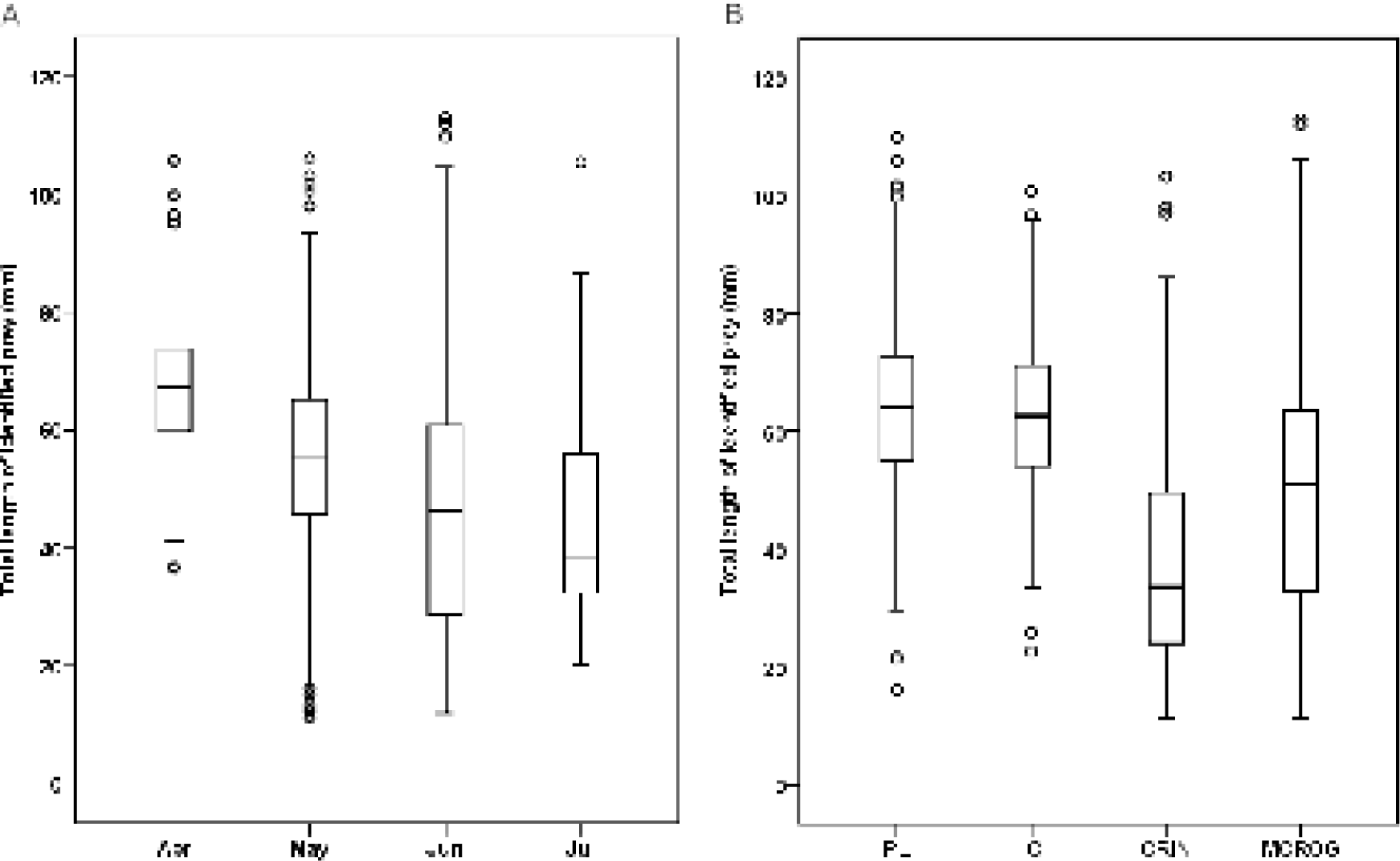
Box graphs show the average total length of the little tern prey on the image analysis. (A) the average total length of prey species compared between months, (B) the average total length of prey species compared between breeding stages. PL, pre-laying; IC, incubation; CRIN, chick-rearing in the nest; MCROG, mobile chick-rearing on the ground.

**Fig 6A, 6B.**
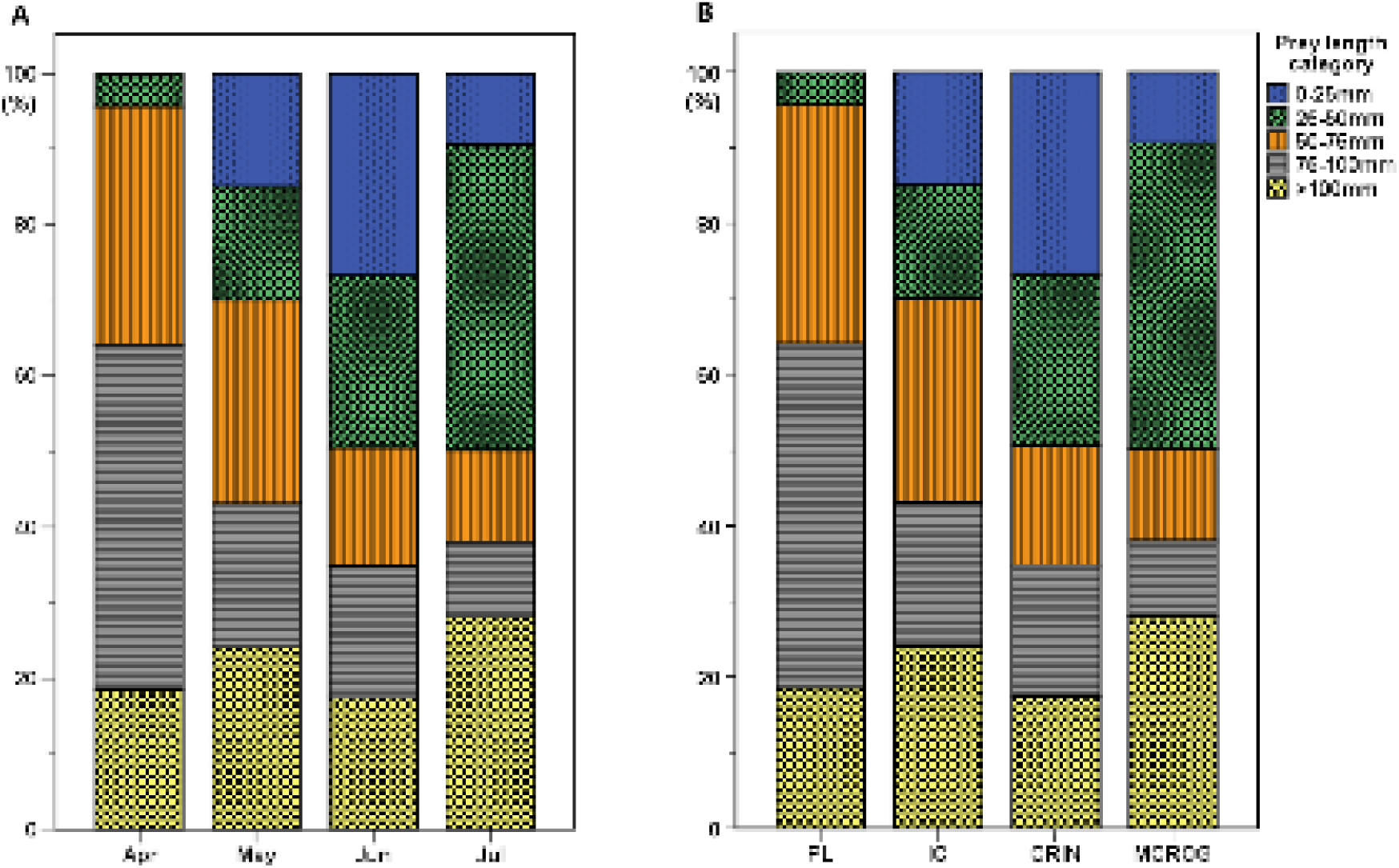
Occurrence frequency of prey length categories of the little tern classified on the images analysis. (A) bar graph of prey length categories compared between months, (B) bar graph of prey length categories compared between breeding stages. PL, pre-laying; IC, incubation; CRIN, chick-rearing in the nest; MCROG, mobile chick-rearing on the ground.

The occurrence frequency of prey length categories also varied significantly among the breeding stages (chi-square test: χ^2^ = 221.217, df = 12, *P* < 0.001). Partitioning of the contingency table and analysis of adjusted residuals indicated that, compared to a chance distribution, 1–25 mm and 50–75 mm categories were overrepresented and underrepresented, respectively, at the chicks-in the nest stage. On the other hand, 50–75 mm category was preferred for the pre-laying and incubation stages, however, where a significant difference was not detected between the both in prey length (Student *t*-test: n = 278, t = 1.055, *P* = 0.292).

Prey length differed significantly between pond smelt and non-pond smelt groups (Mann-Whitney U-test: n = 962, z = –16.012, p < 0.001), where the average of the former showed approximately twice higher value than that of the latter (56.48 ± 18.69 mm, n = 783, range = 11.49–112.93 mm; 29.29 ± 12.51 mm, n = 179, range = 11.27–66.44 mm, respectively). It was pond smelt for little terns to forage most frequently throughout the breeding season, thus showing its greatest prey availability. The length of pond smelt used as prey also differed significantly among 4 breeding stages and months (Kruskal-Willis test: n = 781, χ^2^ = 118.250, *P* < 0.001; n = 981, χ^2^ = 104.624, p < 0.001; Fig 7A and 7B). The medium and larger pond smelts >50 mm and the small ones <5 mm were delivered for females on the mating period and young in the nest, respectively.

**Fig 7A, 7B.**
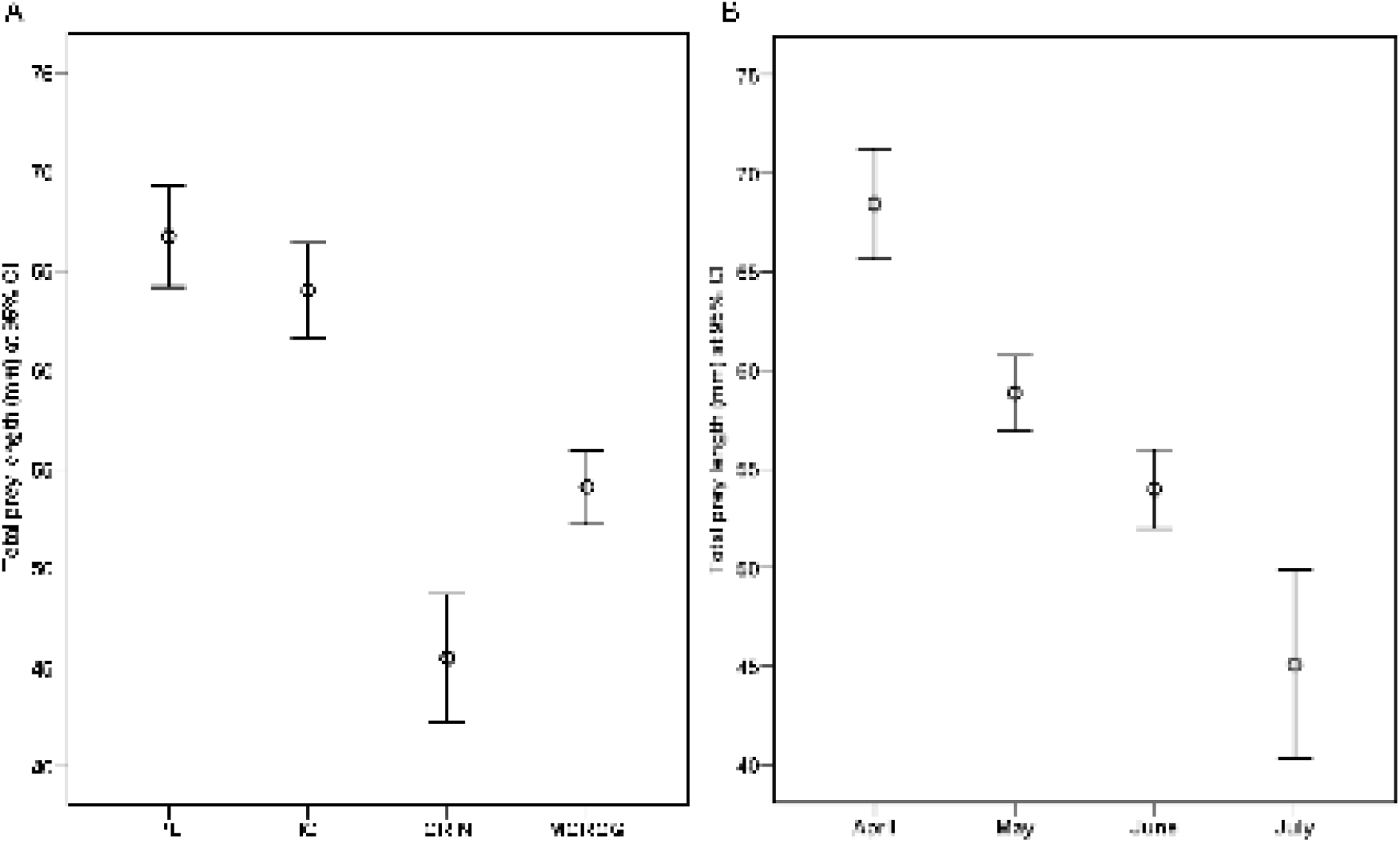
Error-bar plots show the average pond smelt length on the little tern prey images. (A) the total length of the identified pond smelt compared at a 95% confidence interval between breeding stages. (B) the total length of the identified pond smelt compared at 95% confidence interval between months. PL, pre-laying; IC, incubation; CRIN, chick-rearing in the nest; MCROG, mobile chick-rearing on the ground.

In terms of the survival condition of pond smelts, both environmental factors were estimated within a normal range during the entire breeding season from 9th of April to 15th of July [Fig 8 and Fig 9]. The average WST and DO were 22.40 ± 4.35°C (range = 8.27–30.01°C) and 9.49 ± 0.98 mg/L (range = 7.22–12.28 mg/L), respectively. However, they showed a contrary result in July when terns flew away. Between DO between the 1st to 15th and 16th to 31th of July, there was not found a significant difference, whereas those in WST did (Mann-Whitney *U*-test: *P* < 3.313, *P* < 0.001, respectively; Table 2). In addition, DO between 15th and 16th of July did not differ, whereas WST did (Mann-Whitney *U*-test: *P* = 0.186, Student *t*-test: *P* = 0.004, respectively; Table 2). Therefore, a threshold WST for little terns looks like between 29.11°C and 30.04°C, the average WST of their presence (15th of July) and non-presence days (16th of July) [Table 2 and Fig 8], respectively.

**Table 2.**
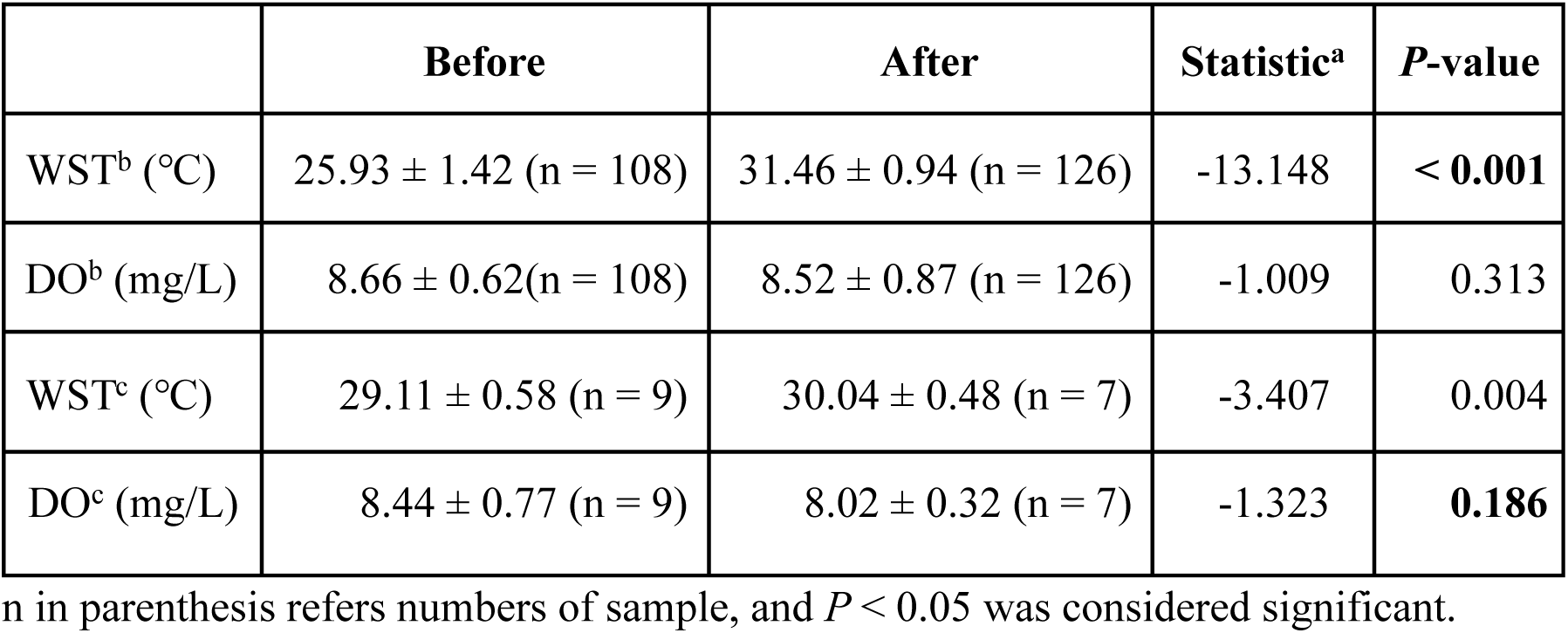
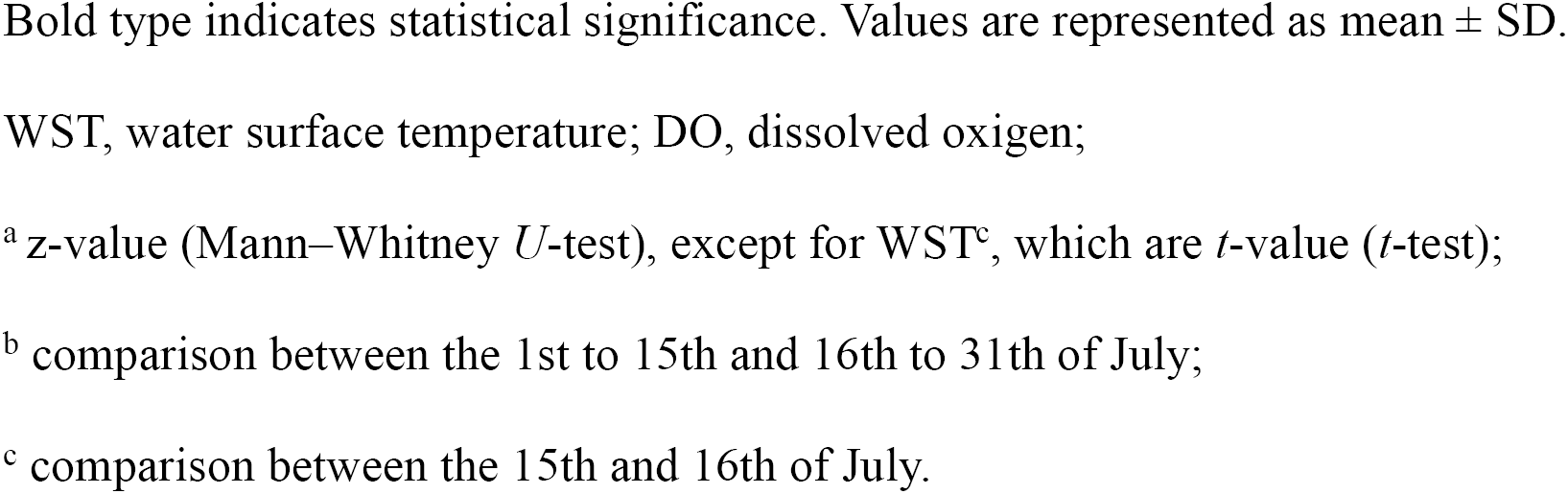
Water factors of the day when little terns flew away analyzed by Mann-Whitney *U* test and *t*-test.

|  | Before | After | Statistic<sup>a</sup> | *P*-value |
| --- | --- | --- | --- | --- |
| WST<sup>b</sup> (°C) | 25.93 ± 1.42 (n = 108) | 31.46 ± 0.94 (n = 126) | -13.148 | < 0.001 |
| DO<sup>b</sup> (mg/L) | 8.66 ± 0.62(n = 108) | 8.52 ± 0.87 (n = 126) | -1.009 | 0.313 |
| WST<sup>c</sup> (°C) | 29.11 ± 0.58 (n = 9) | 30.04 ± 0.48 (n = 7) | -3.407 | 0.004 |
| DO<sup>c</sup> (mg/L) | 8.44 ± 0.77 (n = 9) | 8.02 ± 0.32 (n = 7) | -1.323 | **0.186** |

n in parenthesis refers numbers of sample, and *P* < 0.05 was considered significant.
Bold type indicates statistical significance. Values are represented as mean ± SD.
WST, water surface temperature; DO, dissolved oxygen;
<sup>a</sup> z-value (Mann–Whitney *U*-test), except for WST<sup>c</sup>, which are *t*-value (*t*-test);
<sup>b</sup> comparison between the 1st to 15th and 16th to 31th of July;
<sup>c</sup> comparison between the 15th and 16th of July.

**Fig 8.**
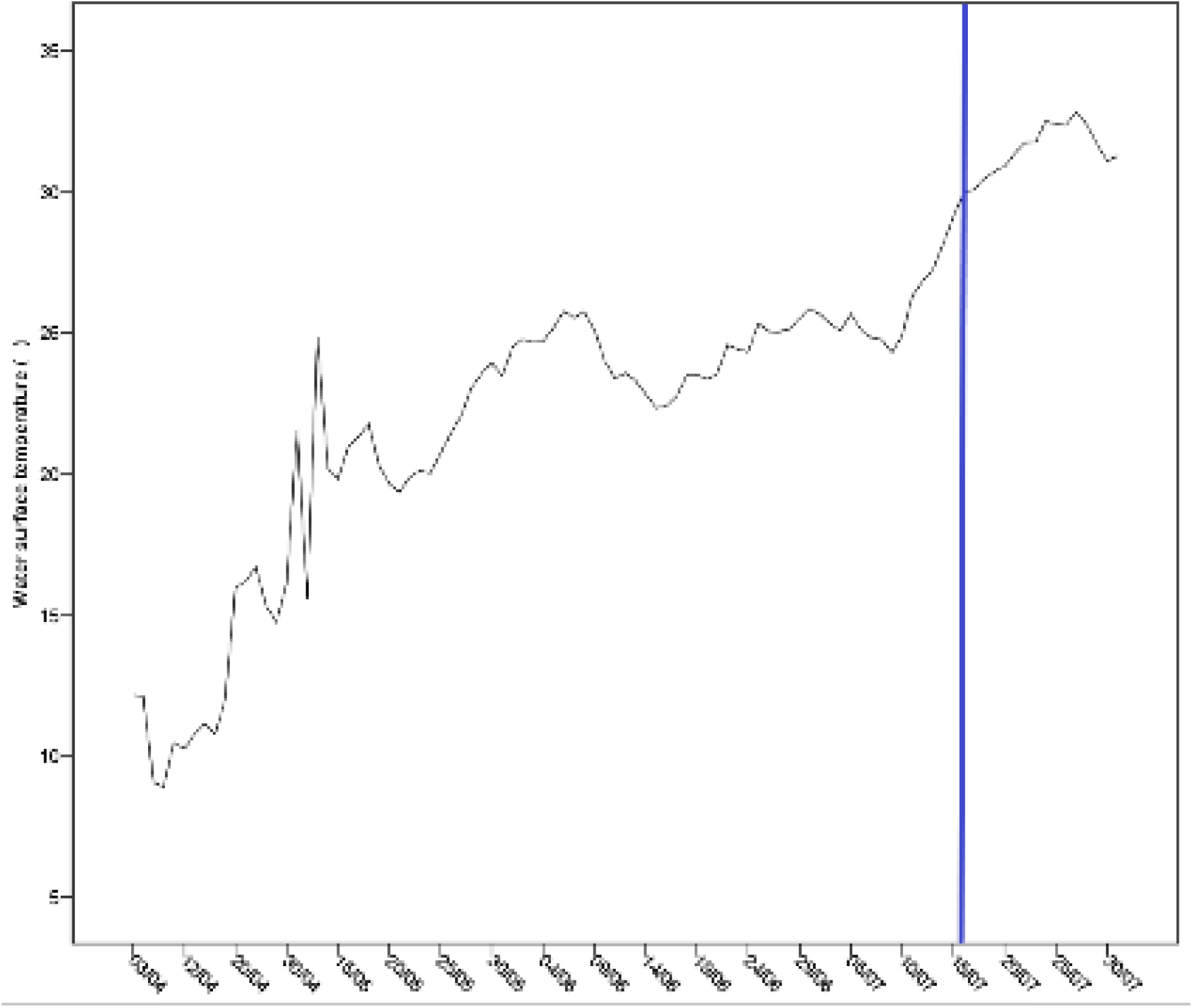
Line graph of the water temperature in Andong Lake of inland Korea during the little tern breeding season. A vertical line in a graph represents the day (16th of July) when the terns flew away for another foraging place.

**Fig 9.**
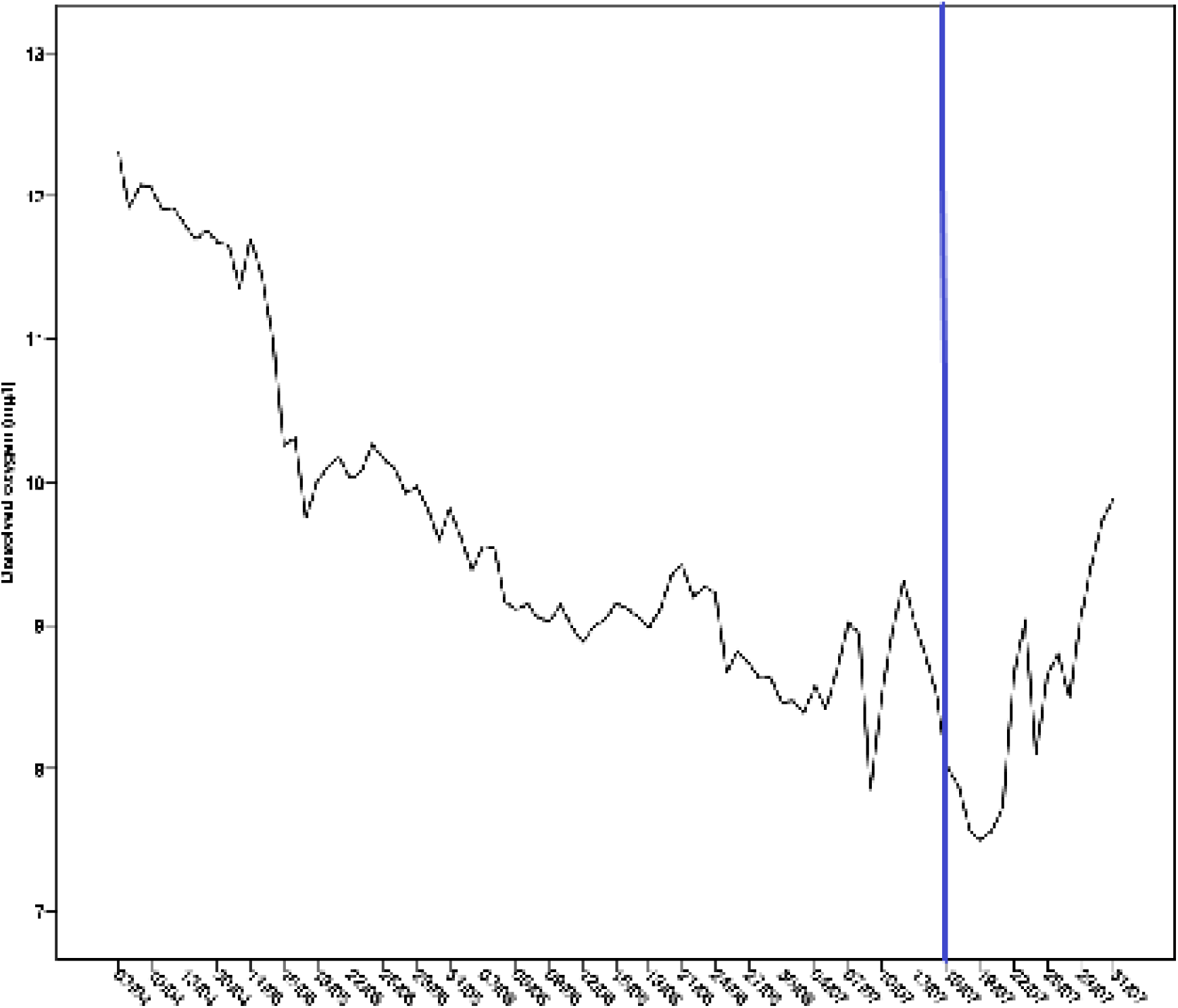
Line graph of the water temperature in Andong Lake of inland Korea during the little tern breeding season. A vertical line in a graph represents the day (16th of July) when the terns flew away for another foraging place.

## Discussion

RCVCs that we set on the islet to capture the images that the little tern had prey in its bill was much effective for identifying their prey species. Non-interference in the breeding activity on colony and revealment of the greater part of prey species were the substantial merit of this method. Above all things, we could record foraging activities at dawn and after sunset attributed to IR mode, which helped for taking all aspects of prey. Besides, the estimation of PTL and LCL on a captured image reflected the actual length approximately. The maximum value of PTLs was 112.93 mm in the current study, which was similar to or did not exceed that of the previous study in Andong Lake (74.0–105.1 mm in standard length) [48].

Eleven fish species including pond smelt and largemouth bass dominated tern’s diet, and insects and crustaceans, also rare in other habitats [3, 5, 27], were not detected, which probably seems to result from the year effect. It was observed in the previous year that a chick fed on a butterfly brought to itself (unpubl. data).

The length of prey delivered to little tern chicks in this study (mean = 45.86 ± 20.55 mm) was similar to that of the UK (30–70 mm) [3]. Generally, the prey type of terns is affected by the fish abundance of foraging place [49, 50]. The ichthyofauna in Andong Lake that pond smelt is abundant [16] provided an excellent opportunity for little terns to forage on it. For the mating season (in particular, April), they preyed mainly on medium-or large-size pond smelts >50 mm congregating for spawning in shallow water [Fig 7A and 7B]. On the contrary, for the chicks in the nest (in particular, July), small-size pond smelts <50 mm were delivered mainly. However, the frequency of very small fries <25 mm was there comparatively rare (16.4%; Fig 7A and 7B). These results explain different prey preference according to breeding stages [11].

Although pond smelts accounted for 58% and 82.2% of the diet by number for the chicks on the nest or mobile ones on the ground, respectively, largemouth bass fries were also consumed as the important prey for both ones (33.6%, 12%, respectively). In particular, when the opportunistic little tern foraged on very small fries <25 mm for chicks, it preferred for bass fries [Fig 10A and 10B], which supports that in early June, its hatching season, very small largemouth bass fries were the most abundant in the water surface. It might be the result of a breeding strategy for being timed to coincide with peak fish abundance [51, 11]. It probably seems that the diet of the seabird in this inland lake reflected variability in food-web composition due to natural or human-induced environmental change [52, 53, 36].

**Fig 10A, 10B.**
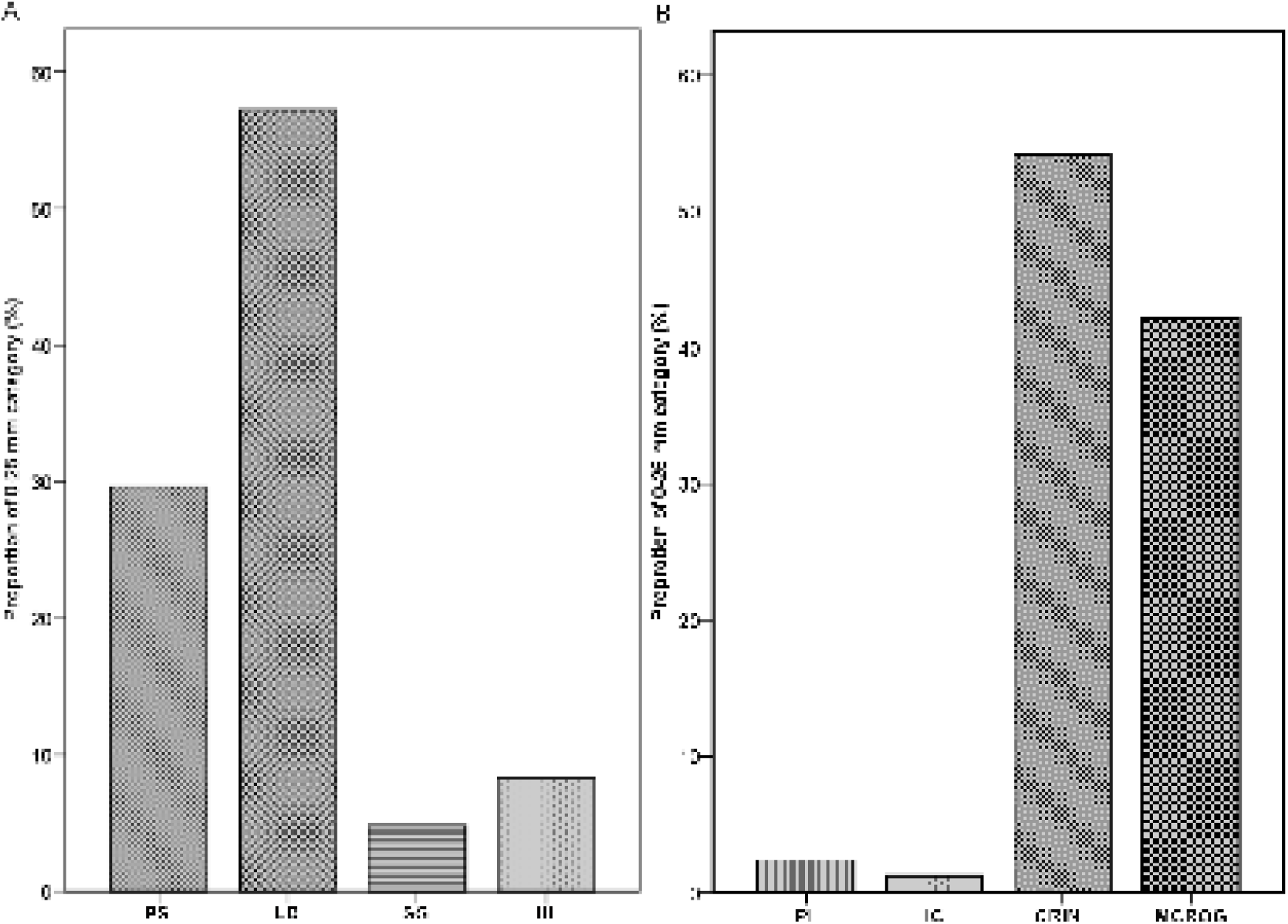
Occurrence frequency of 0-25 mm category of little tern preys classified on the images analysis. (A) bar graph of 0-25 mm prey length category compared between, (B) bar graph of 0-25 mm prey length category compared between breeding stages. PS, Pond Smelt; LB, Largemouth bass (*Micropterus salmoides*); SG, Skygager (*Erythroculter erythropterus*); UI, unidentified; PL, pre-laying; IC, incubation; CRIN, chick-rearing in the nest; MCROG, mobile chick-rearing on the ground.

What is most interesting is that cold-water pond smelts were taken in the hot summer by terns. The DO of lake water has a strong influence on the population dynamics of the pond smelt, which is almost absent in anoxic areas where it is below 3mg/l [54, 55]. However, it was much above 3mg/l in this lake throughout the breeding period [Table 2 and Fig 9], which did not thus influence on the pond smelt population and tern’s foraging activity. The cold-water pond smelt is commonly a eurythermal and euryhaline fish with high adaptability in various environmental factors such as turbidity, salinity, and temperature and can thus survive in water of low quality [56–58]. In the present study, it survived between WST 29.11 and 30.04°C [Table 2], which indicates that a threshold value of WST for pond smelts was statistically 29.58 °C on average. These results support the prey abundance hypothesis that, when cold-water pond smelts might wholly swim down into the deeper lake in the hot summer [20, 21], the terns might also leave their colony for another foraging place with higher prey availability. Besides, results in this study were similar to suggestions that their lethal survival temperature was 29.1°C in the experimental condition [57, 59].

Although the pond smelt was human-introduced species, the predatory tern, originally a seabird, utilized the landlocked fish effectively in the lake. Since it prevented the excessive increase of the population of exotic largemouth basses, conserving the rare breeding colony in this inland lake could largely contribute to maintaining the health of the lake ecosystem and biodiversity. By the way, there may come in the future a day when the WST exceeds the critical temperature in the lake due to unexpected dramatic climate change before little tern’s breeding is finished, and thus mass mortality of pond smelts may occur, such as in Lake Kasumigaura [59], Japan, and Woon am Reservoir, Korea [60]. Therefore, multi-annual diet research is required for a deeper understanding of its prey fluctuation and further measures for habitat conservation.

## Acknowledgements

This work was supported by National Research Foundation of Korea(NRF) grant funded by the Korea government(MSIT) (2013M3A9A5047052 and 2017M3A9A5048999). In addition, it was conducted with the production of a documentary (The secret of the little tern seabird of the inland lake; aired on 25 October 2018; KBS1), which was approved by Andong City and K-water (Permit: K-water Andong 2018-8). We wish to thank Dong-Man Shin, director; Sang-Sup Yeom, co-director; and Sung-Il Shin and Woo-Kyung Lee, cinemaphotopgraphers, that cooperated to collect image data.

